# LncRNA-536 and RNA Binding Protein RBM25 Interactions in Pulmonary Arterial Hypertension

**DOI:** 10.1101/2024.08.27.610011

**Authors:** Aatish Mahajan, Ashok Kumar, Ling Chen, Navneet K. Dhillon

## Abstract

**OBJECTIVE:** Hyperproliferation of pulmonary artery smooth muscle cells (PASMCs) is one of the essential features of the maladaptive inward remodeling of the pulmonary arteries in pulmonary arterial hypertension (PAH). In this study, we define the mechanistic association between long-noncoding RNA: ENST00000495536 (Lnc-536) and anti-proliferative HOXB13 in mediating smooth muscle hyperplasia.

**METHODS:** Antisense oligonucleotide-based GapmeRs or plasmid overexpressing lnc-536 were used to evaluate the role of lnc-536 in mediating hyperproliferation of PDGF-treated or idiopathic PAH (IPAH) PASMCs. Further, we pulled down lnc536 to identify the proteins directly interacting with lnc536. The in-vivo role of lnc-536 was determined in Sugen-hypoxia and HIV-transgenic pulmonary hypertensive rats.

**RESULTS:** Increased levels of lnc-536 in PDGF-treated or IPAH PASMCs promote hyperproliferative phenotype by downregulating the HOXB13 expression. Knockdown of lnc-536 *in-vivo* prevented increased RVSP, Fulton Index, and pulmonary vascular remodeling in Sugen-Hypoxia rats. The lncRNA-536 pull-down assay demonstrated the interactions of RNA binding protein: RBM25 with SFPQ, a transcriptional regulator that has a binding motif on HOXB13 exon Further, The RNA-IP experiment using the SFPQ antibody showed direct interaction of RBM25 with SFPQ and knockdown of RBM25 resulted in increased interactions of SFPQ and HOXB13 mRNA while attenuating PASMC proliferation. Finally, we examined the role of lnc-536 and HOXB13 axis in the PASMCs exposed to the dual hit of HIV and a stimulant: cocaine as well.

**CONCLUSION:** lnc-536 acts as a decoy for RBM25, which in turn sequesters SFPQ, leading to the decrease in HOXB13 expression and hyperproliferation of smooth muscle cells associated with PAH development.

## INTRODUCTION

Pulmonary arterial hypertension (PAH) is a heterogeneous and fatal disease characterized by pulmonary vascular remodeling ^1,2^ right ventricular failure (RVF), and ultimately, death ^3,4^. Available PAH treatments bring relief from symptoms but can’t cure the disease ^5^. Therefore, a need exists to identify novel targets that drive pulmonary vascular remodeling to design alternative therapeutic approaches.

Long non-coding RNAs (lncRNAs), a novel class of non-coding RNAs was recently discovered with the advancement of RNA-sequencing techniques^6^. LncRNAs are longer than 200bp and do not have protein-coding ability^7^ however they can activate or silence their target genes^7^. Some lncRNAs function as decoys, guides, or scaffolds to a diverse set of transcription regulators while others can function as miRNA sponges therefore regulating the levels and downstream functions of mRNAs^7^. Dysregulation of lncRNAs has been implicated in the development of many human diseases such as cancers and neurological disorders^8,9^, however, only a few studies have demonstrated their role in pulmonary hypertension^10–12^.

Pulmonary artery smooth muscle cells (PASMCs) are one of the two most studied vascular cell types in the attempt to delineate the mechanism(s) responsible for PAH ^13^. In our previous study we reported significant alterations in the levels of various lncRNAs in the hyperproliferative PASMCs ^14^. Based on the fold change and raw intensity in the microarray analysis, ENST00000495536 (ENST-536/ lnc-536) was identified as one of the top up-regulated vascular disease-related lncRNA. Lnc-536 is an intergenic lncRNA located around 12 kbp away from the tumor suppressor gene HOXB13 on the opposite antisense strand (+) of chromosome 17 ^15,16^ . Given that lncRNAs are known to regulate the expression of proximal protein-coding genes in *cis-regulatory* mechanisms,^17^ the gene proximity analysis suggests an association of lncRNA ENST-536 with HOXB13 Therefore, in this study we examined the mechanistic interactions of lnc-536 with anti-proliferative HOXB13 and inquired if these interactions play a role in smooth muscle hyperplasia and PAH, using in vitro, ex vivo, and in-vivo preclinical-models-.

## METHODS

### Human and rat cell cultures

Primary human pulmonary arterial smooth muscle cells (HPASMC) were grown in smooth muscle cell media (SMCM) (Sciencell, USA) with growth factors, 2% fetal bovine serum, and penicillin/streptomycin until 70% confluency on 6 well plates and made quiescent for 48 h with serum-free media and then treated with either PDGF-BB (100ng/ml) for 24h or cocaine (C) (1 µM) and HIV-Tat protein (25 ng/mL) for 48 h based on our previous findings ^18,19^ . IPAH and familial PAH (FPAH) cells were grown in VascuLife® SMC Medium. After appropriate treatments, cells were harvested in TriZol reagent (Invitrogen), and isolation of total RNA and quantitative RT-PCR was performed as described previously ^14^. The qRT-PCR primers for the selected mRNA and lncRNAs were either custom-designed using the PrimerQuest tool and obtained from Integrated DNA technologies or ordered from Qiagen (List of primers used in the study Supplementary Table 1).

Rat pulmonary arterial smooth muscle cells (RPASMCs) were isolated from pulmonary arteries as in our previous published findings ^19,20^. The cells were cultured on a 6-well plate in rat smooth muscle cell media (R-SMCM) (Cell Applications, USA) and used for further experiments.

### Oligonucleotides and plasmid transfections

Antisense-locked nucleic acid (LNA) GapmeRs against lnc536 were designed using Qiagen, a GapmeR designer tool for in-vivo and in-vitro experiments. The top-ranked antisense GapmeRs were ordered along with positive and negative controls. HPASMC were reverse transfected while seeding in 6 well plates at 2 × 10^5^ cells/well with antisense LNA-GapmeR against lnc-536 (LNA-AS536) or scrambled LNA (LNA-Scr) at 40 nM using HiPerfect transfection reagent (Cat#301704; Qiagen). ^14^ After 24 h of transfection, the cells were serum starved for 24 h and then treated with PDGF for 24h or with Cocaine and Tat (C+T) for 48 h. RNA isolation and qRT-PCR were performed to evaluate the knockdown of specific lncRNAs. For RBM25 knockdown in the cells, RBM25 PSI-H1 shRNA (Catalogue # HSH108418-CH1) and shRNA scrambled (Catalogue # CSHCTR001-CH1) clones were obtained from GeneCopoeia Inc and transfected using the manufacturer’s instructions.

The following plasmids were used: customized puc57-Lnc536 (Lifeasible INC), pCMV6-HOXB13-MYC-DDK (RC209991, Origene), and pCMV6-SFPQ-MYC-DDK (RC208454, Origene). To clone the region consisting of Lnc536 into puc57, the region was PCR amplified and inserted into the vector digested with HindIII/ EcoRI followed by ligation. The insertion of lnc-536 was verified by sequencing. HPASMCs were transfected with these plasmids with GeneJuice (Sigma) according to the manufacturer’s protocol. For the negative controls, empty PUC57, and empty pCMV6 were used.

### Proliferation and Apoptosis Assays

The effects of lnc536 knockdown, RBM25 knockdown, or HOXB13 over-expression in cells over-expressing lnc-536 or in cells treated with PDGF-BB or cocaine and Tat (C+T) on the proliferation and apoptosis of control or IPAH HPASMCs were assessed. The assays were performed using CellTiter 96® AQueous One Solution Cell Proliferation Assay (MTS) (Promega, G3582), CyQUANT® Cell Proliferation Assay Kit (Invitrogen, C7026), and edU (5-bromo-2’-deoxyuridine) incorporation assay kit according to the manufacturer’s instructions. ^21^

### Lnc-536 RNA pulldown assay and peptide purification for mass spectrometry

The RNA pulldown assay was performed commercially by Lifeasible INC. First, to obtain lnc-536 RNA the puc57-lnc536 or control puc57 plasmids were linearized with SmaI (Thermo Fisher). After precipitation and purification of linearized DNA, DNA was *in vitro* transcribed according to the manufacturer’s protocol with T7 Phage RNA Polymerase (NEB), and DNA was digested with RQ DNase I (Promega). The remaining RNA was purified with the RNeasy Mini Kit (Qiagen) and biotinylated at the 3’end with the Pierce RNA 3’end biotinylation kit (Thermo Fisher). Both antisense and sense strands of the DNA sequence were amplified to obtain a DNA template for *in-vitro* transcription (Supplementary Figure. E2B) and an antisense DNA template was used as a negative control. For proper RNA secondary structure formation, 150ng of 3’end biotinylated lnc536 RNA or control RNA was heated for 2 min at 90°C in RNA folding buffer (10 mM Tris pH 7.0, 0.1 M KCl, 10 mM MgCl2), and then put on RT for 20 min. Streptavidin magnetic beads were added to the transcribed RNA and incubated for 1h. The magnetic beads and RNA were removed with a magnetic separator.

This labeled immobilized lncRNA was then incubated with the HPASMC protein extract (from 4×10^7^ cells) followed by elution of bound proteins and analysis using liquid chromatography-mass spectrometry (LC-MS). Part of the eluted protein was loaded on the SDS-PAGE for silver staining that showed multiple proteins with the sense RNA strand pull-down (experimental group) whereas the antisense lane (negative control) was almost clear (Supplementary Figure. E2 D). To reduce the complexity of mass spectrometric measurements, samples were eluted stepwise from the beads. The first elution was performed with 5 M Urea, 2 mM DTT, and 50 mM Tris pH 7.5, for 15 min. Thiols were alkylated with 57 mM chloroacetamide for 20 min at RT in the dark. Beads were sedimented and the supernatant was transferred to Microcon YM-30 (Millipore) spin filters (30 kDa cut-off) and centrifuged for 10 min. Proteins from beads and filters were digested overnight with 1 µg LysC (Wako) in 100 µL 2 M Urea, 50 mM Tris pH 7.5, and combined. Beads were washed with 100 µL ABC and supernatant was transferred to the digestion and the mixture was digested with 1 µg Trypsin (Promega) overnight at RT. The remaining proteins on beads were eluted for 10 min at 95°C in 4% SDS, 50 mM HEPES pH 7.6, 100 mM NaCl, and 10 mM DTT. Subsequently, the standard protocol of filter-aided sample preparation (FASP) was performed18 including RNase T1 (1000 U/µL, Thermo Scientific) and RNase A (4 µg/µL, Thermo Scientific) digestion using 1 µL each to elute crosslinked peptides. Peptides from both fractions were acidified by trifluoroacetic acid (TFA) to a final concentration of 0.1% and purified on multi-stop-and-go tips (StageTips) containing a stack of three C18-disks19. Peptides were resolved in 1% acetonitrile and 0.5% formic acid and further analyzed by liquid chromatography/mass spectrometry (LC/MS).

### RNA-Immunoprecipitation

After appropriate treatments, PASMCs were scraped and resuspended in lysis buffer (50 mM Tris-HCl pH 7.4, 100 mM NaCl, 1% NP-40, 0.1% SDS, 0.5% Sodium deoxycholate and protease inhibitors) for 10 min. Five percent of lysate was taken as an input sample. Protein A agarose beads (Invitrogen) (50 μL) were incubated with 4 μg of RBM25 or SFPQ antibody (Abcam) for 1 h at room temperature. Antibody-coupled beads were then incubated with the lysate for 1 h at 4 °C. After washing the beads three times for 10 min in high salt buffer (50 mM Tris-HCl, pH 7.4, 1 M NaCl, 1 mM EDTA, 0.1% SDS, 0.5% Sodium deoxycholate, 1% NP-40), RNA was extracted using Trizol and chloroform, reverse transcribed and amplified using qPCR^22^

### Animal experiments

To see if inhibiting Lnc-536 could prevent pulmonary vascular dysfunction we used Sugen-hypoxia and HIV-transgenic (Tg) rat PH models. Sprague Dawley male rats subcutaneously treated with 20 mg/kg body wt. of Sugen 5416 ^23^ (day 0) and exposed to 10% oxygen environment for 3 weeks were injected LNA-AS536 GapmeRs through tail vein at a dose of 5mg/kg at day 4,9,14. Rats treated with LNA-Scr GapmeRs at a dose similar to LNA-AS536 served as controls ^24–26^) . To see the in-vivo role of lnc-536 in HIV-Tg rats, 8 to 9-month-old male HIV-Tg Fisher rats exposed to cocaine (40mg/kg IP, daily ) for 21 days were treated with LNA_Scr or LNA-AS536 on days 4, 9, and 14. Transthoracic echocardiography was performed on all rats anesthetized with Ketamine/Xylazine mixture (80 mg/kg:10 mg/kg, IP), on days 0and 21 to measure the end-diastolic area (EDA) and ejection fraction (EF). At the end of treatments on day 21, echocardiography was followed by catheterization of anesthetized rats for hemodynamic analysis. Finally, tissues were harvested and the RV/LV + Septum ratio (Fulton Index) was calculated to assess the RV hypertrophy (RVH) as described in our previous published findings ^27^) . The efficiency of LNC536 knockdown in PASMCs from AS-536 treated rats was confirmed using real-time PCR. The animals were housed at the University of Kansas Medical Center (Kansas City, KS) in strict accordance with the National Institutes of Health (NIH) and Institutional Animal Care and Use Committee (IACUC) guidelines (Protocol Number: 2021-2067).

### Statistical Analysis

Data were analyzed using Prism 10 software (GraphPad Sofware Inc.) and displayed in mean ± SD. Pairwise comparisons between experimental and control groups were made using paired or unpaired 2-tailed Student’s t-test as appropriate. One-way analysis of variance (ANOVA) was used to compare two groups of continuous variables with normal distribution, and two-way ANOVA, followed by post hoc tests, was used to compare multiple group comparisons as previously described ^14^. Unless otherwise stated, P < 0.05 was considered statistically significant.

## RESULTS

### Lnc-536 promotes smooth muscle proliferation by attenuating anti-proliferative HOXB13 expression

To study the functional relevance of lnc536 in smooth muscle cells we first examined the proliferation of HPASMCs over expressing lnc-536. Cells transfected with puc57-Lnc536 having higher levels of lnc-536 resulted in the augmentation of smooth muscle proliferation as shown in Supplementary Figure E1A (i-iii). Similarly, HPASMCs exposed to PDGF mitogen exhibiting hyperproliferative phenotype had higher expression of lnc-536 (Figure 1A i-iii). Further, the knockdown of lnc536 using antisense lnc-536 GapmeRs (LNA-AS536) resulted in a decrease in the PDGF-mediated smooth muscle hyper-proliferation as shown by MTT, Cyquant and EdU incorporation proliferation assays (Figure 1B ii-iv).

**Figure 1.**
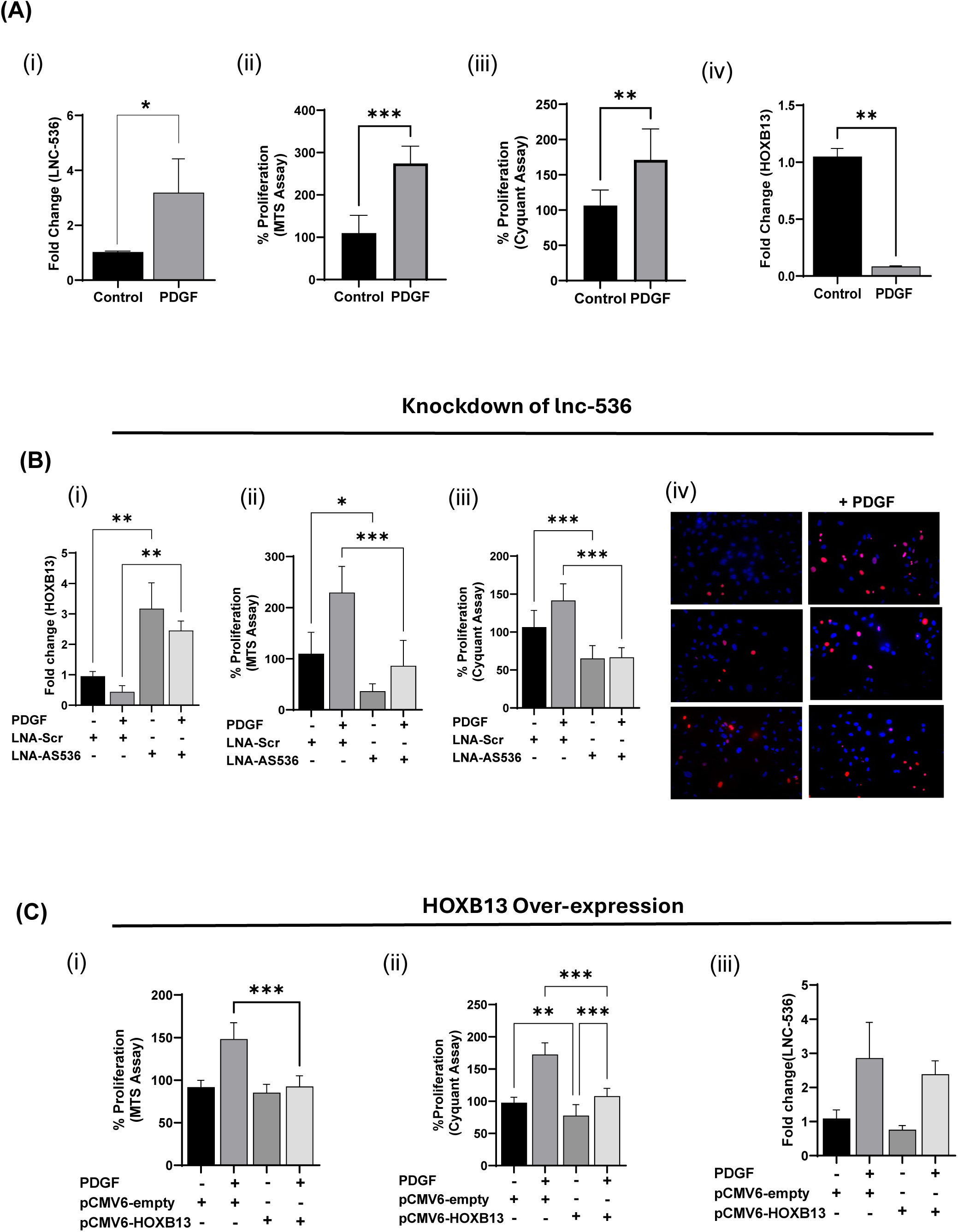
(A) Effect of PDGF treatment on the expression of lncRNA536 (i), cell proliferation as assayed by MTS (ii) and Cyquant (iii) and on HOXB13 levels (iv) in human pulmonary arterial smooth muscle cells (HPASMCs). *p<0.05, **p <0.01, *** p<0.001 vs control (B). Effect of lncRNA knockdown in PDGF-treated HPASMCs on HOXB13 expression (i) and cell proliferation/survival as assayed by MTS (ii) , Cyquant (iii), and Edu (iv). HPASMCs were transfected with LNA-gapmeR against lnc-536 (lnc536-AS1,40nM) or LNA-gapmeR scrambled control (n=3) for 24h followed by 24 h PDGF (100ng/ml) treatment. **p <0.01, *** p<0.001. (C). Effect of HOXB13 over-expression in PDGF treated HPASMCs on cell proliferation as assayed by MTS (i), Cyquant (ii), and effect on lnc-536 expression (iii). HPASMC were transfected with 2µg of plasmid over-expressing HOXB13 (pCMv6-HOXB13), or pCMv6-empty as control (n=3) for 24h followed by treatment with PDGF (100ng/ml) for 24h. **p <0.01, *** p<0.001. Data is an average of a minimum of three independent experiments performed in triplicates. Representative images of Edu staining are shown.

Further, HPASMCs transfected with plasmid over-expressing lnc536 (Supplementary Figure E1A iv) and PASMCs treated with PDGF led to a decrease in the expression of tumor suppressor HOXB13 gene (Figure 1A iv). Alternatively, the knock-down of lnc536 in HPASMCs (Figure 1Bi) treated with or without PDGF resulted in the increased expression of HOXB13 with the corresponding significant decrease in cell proliferation compared to cells transfected with scrambled control. These results show that silencing lncRNA-536 results in an anti-proliferative phenotype by increasing the expression of HOXB13.

To confirm the direct effect of HOXB13 on HPASMC phenotype, we transfected cells with the plasmid over-expressing HOXB13 (pCMV6-HOXB-13) in the presence and absence of lnc-536 overexpression (PUC57-lnc536) or PDGF treatment. Indeed, the overexpression of HOXB13 reduced the lnc-536 mediated hyper-proliferation of HPAMCs (pCMV6 -empty+ PUC57 lnc536 vs pCMV6-HOXB13 + PUC57 lnc536) (p<0.001) as evident from MTT, Cyquant and Edu assays (Supplementary Figure E1B i-iii). Further, the cell proliferation was also found to be significantly decreased in cells only over-expressing HOXB13 (pCMV6-empty vs pCMV6-HOXB13) as assayed by Cyquant assay (Supplementary Figure E1Bii). Similarly, HOXB13 overexpression could significantly attenuate the PDGF-mediated proliferative phenotype of HPASMCs (pCMV6-empty+ PDGF vs pCMV6-HOXB13 + PDGF) as shown in Figure 1C (i-ii). Correspondingly as shown in Supplementary Figure E1B(iv), no significant difference in lnc536 expression was observed when the HPASMCs were transfected with plasmid over-expressing HOXB13 (pCMV empty vs pCMV-HOXB13), whereas over-expression of HOXB13 in cells overexpressing lnc536 resulted in significant decrease in the expression of lnc536 (pCMV6 empty + PUC57-lnc536 vs pCMV6-HOXB13 + PUC57-lnc536), thereby suggesting feed-forward regulation of lnc-536 by downstream HOXB13. However, no significant decrease in the lnc536 expression was observed in PDGF-treated cells over-expressing HOXB13 compared to PDGF-treated control cells (Figure 1C iii).

### Knockdown of lnc-536 attenuated pulmonary vascular remodeling and right ventricular dysfunction in Sugen-Hypoxia PAH rats

Conservation analysis of lnc536 using the Ensemble genome browser and MView tool indicated conserved regions across all vertebrate species including rats (Ensemble genome browser 96; human genome version, GRCh37.p13) as shown in (Supplementary Figure E2A). So, to determine the in-vivo role of lncRNA-536 in the development of PAH, we exposed rats to hypoxia for 21 days after one Sugen injection in the presence of either treatment with LNA-Scr or LNA-AS536. Echocardiography analyses observed an increase in normalized Ejection Fraction (ΔRV/LV EF) and a reduction in normalized End Diastolic Area (ΔRV/LV EDA) in Su-Hypoxia PAH rats treated with LNA_AS-536 treatment compared to Su-hypoxia rats treated with scrambled control (Figure 2A). The ratio of RV acceleration time (AT) and ejection time (ET) was also found to be significantly low in LNA-AS-536 treated rats. Right ventricular systolic pressure (RVSP) and Fulton Index, were significantly decreased in the Sugen-hypoxia rats on treatment with LNA-AS536 compared to Sugen -hypoxia rats treated with LNA-Scr GapmeRs (Figure 2A),. Correspondingly, less RV collagen deposition (Figure 2B) and reduced pulmonary vascular remodeling (Figure 2C) were observed in Sugen-hypoxia rats exposed to LNA-AS536 compared to scrambled control. Analysis of lncRNA-expression levels in pulmonary arterial smooth muscle cells isolated from rats (RPASMCs) indicated a decrease in lnc-536 in Sugen-hypoxia rats treated with AS-536 that resulted in augmentation of HOXB13 expression (Figure 2D (i-ii). Further, the silencing of lnc-536 in Sugen-hypoxia rats resulted in attenuation of RPASMCs cell viability and cell proliferation as measured by MTS and Cyquant assays compared to rats treated with scrambled GapmeRs, which was further confirmed by EdU incorporation assay (Figure 2 E (i-iii).

**Figure 2.**
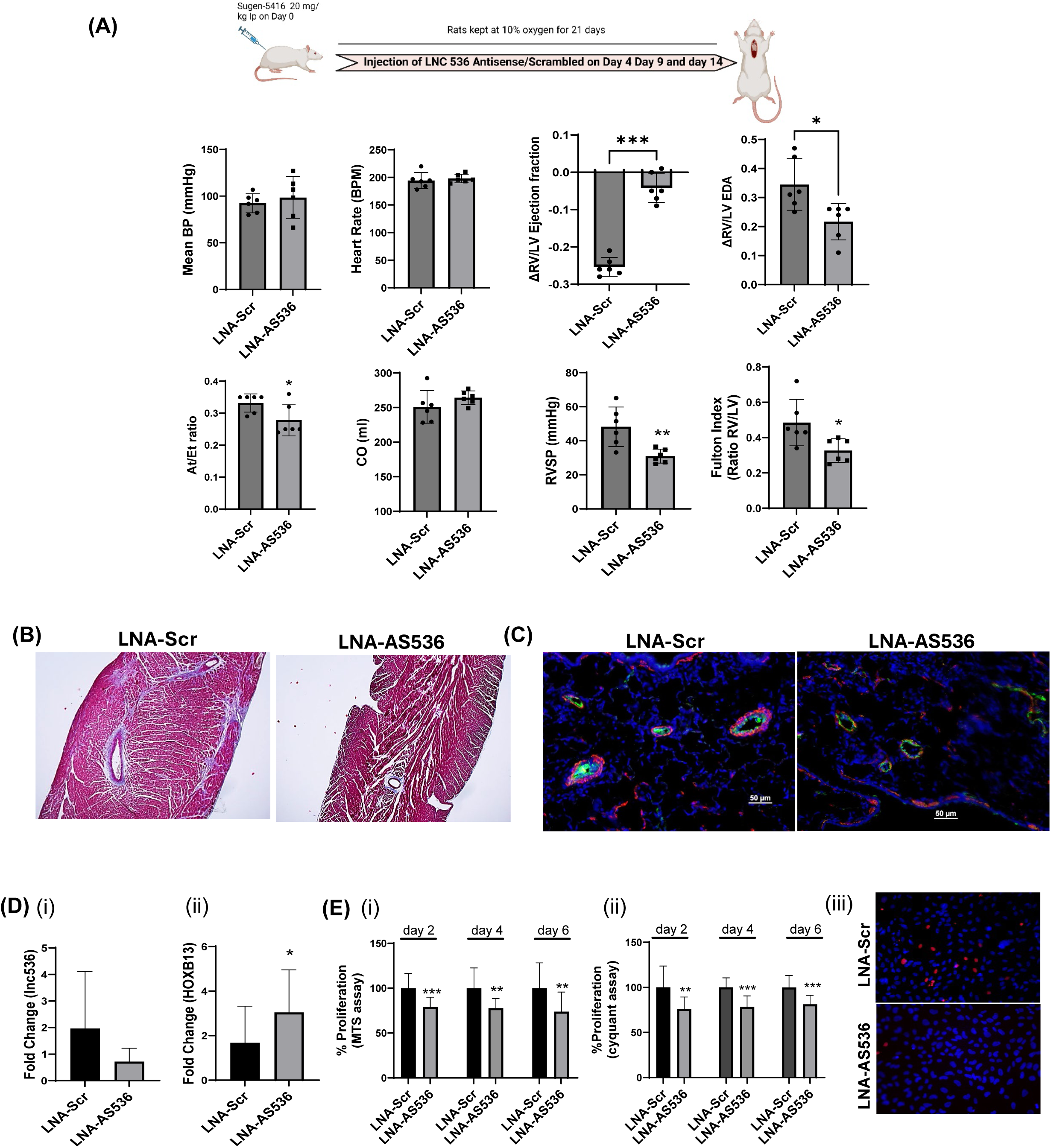
Effect of knockdown of lnc536 in Sugen-hypoxia PAH rats. Adult male fisher rats received Su5416 (20 mg/kg) followed by immediate hypoxia exposure ( 10% O_2_ for 21 days). Rats (n=6/group) were treated with LNA-gapmeR against lnc-536 (LNA-AS536) or LNA-gapmeR scrambled control (LNA-Scr) ( 5 mg/kg body wt.) on days 4, 9, and 14. (A) Echocardiography and hemodynamic analysis: Mean BP, Heart rate, ΔEV/LV Ejection fraction(EF), ΔRV/LV end-diastolic area (EDA), At/Et, cardiac output (CO) along with RVSP, Fulton index data are shown. Representative micrographs of trichrome stained right ventricles (B) and alpha-smooth muscle actin (red) and vWF ( green) immuno-fluorescence stained lungs (C) from Sugen-hypoxia rats treated with LNA-AS536 or scrambled control (D) Expression of lnc536 (i) and HOXB13 (ii) in pulmonary arterial smooth muscle cells isolated from rats treated with LNA-AS536 or LNA-Scr control (24h). (E) Proliferation analysis of PASMCs isolated from rats were grown under no serum conditions for 2, 4 or 6 days assayed using MTS (i) and Cyquant (ii). Representative images of Edu (iii) stained RPASMCs serum starved for 2 days. *p<0.05, **p <0.01, *** p<0.001 vs LNA gapmeR scrambled control.

### Lnc536 Acts as a Decoy for RNA-binding protein RBM25

#### Analysis of proteins directly associated with lnc536

To further obtain insight into the mode of action of the ENST-536 lncRNA in regulating the smooth muscle phenotype, an *in-vitro* lncRNA pull-down assay was performed to identify the RNA-protein interactions (Supplementary Figure E2B-C). Mass spectrometry analysis of RNA-bound proteins detected 196 proteins bound to the lnc-536 sense strand and 89 proteins in the anti-sense negative control. The 119 proteins identified specifically only in the sense strand group were then ranked based on Unused ProtScore, a true indicator of protein confidence (when Unused >2.0, the confidence for detected protein is above 99%) and analyzed using various bioinformatics tools. KEEG pathway annotation and analysis identified spliceosome, ribosome/ribosome biogenesis, and RNA transport as some of the top 10 pathways associated with sense-strand pulled-down proteins. RNA binding proteins (RBP) that form ribonucleoprotein(s) (RNP)complexes were identified as one of the major groups of pulled-down proteins by lnc536 and that included RNA binding motif 25 (RBM25). RBM25 was among the top 10 specific proteins that were found bound to lnc536 and had an Unused score of >10 (Supplementary Figure E2E). It is a highly conserved RNA binding protein that acts as a splicing regulator by interacting with multiple splicing factors and spliced mRNAs ^28^.

#### RNA-immunoprecipitation (RNA-IP) using RBM25 antibody

To validate the identified interactions between RBM25 and lnc536, we confirmed the mass spectrometry results by RNA immuno-precipitation using an anti-RBM25 antibody. Firstly, we performed RNA pull-down using RBM25 antibody on cells transfecting with and without PUC57-lnc536 overexpressing plasmid. RT-PCR analysis confirmed that the amount of lnc536 was significantly high in the RNA-RBM25 pulled-down complex from the cells overexpressing lnc536 compared to the cells transfected with the empty plasmid control and in parallel the levels of HOXB13 in the complex were significantly low (Figure 3A i-ii). Similarly, RNA-IP on HPASMCs treated with PDGF confirmed the interactions of lnc536 and HOXB13 via RBM25 as observed by higher levels of lnc536 and lower levels of HOXB13 pulled down by RBM-25 antibody (Figure 3B i-ii).

**Figure 3.**
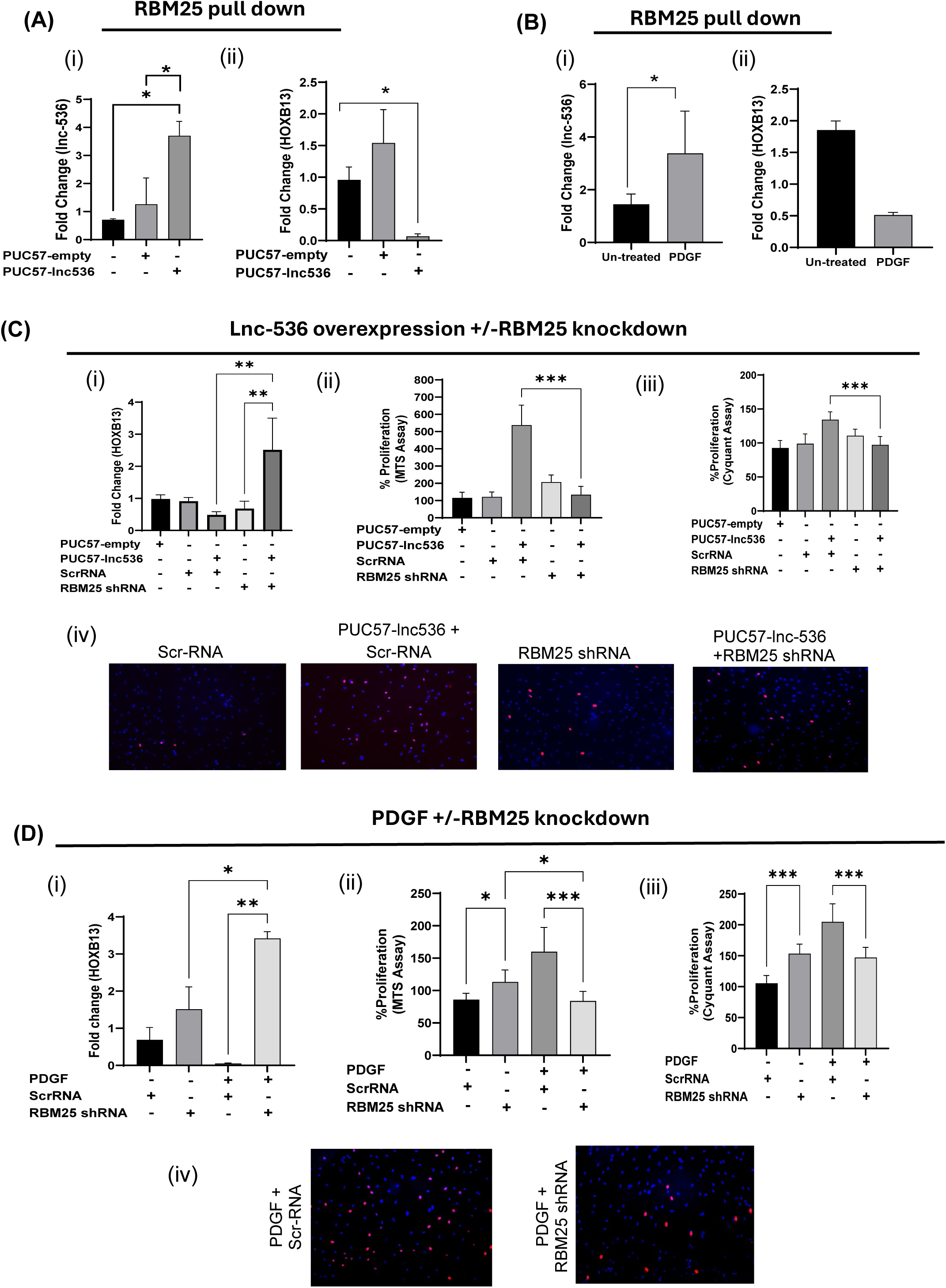
(A) RBM25-Immunoprecipitation (IP) in HPASMCs transfected with and without plasmid over-expressing lnc536. Levels of LNC536 (i) and HOXB13 (ii) RNA after incubation of total cellular extract with RBM-25 antibody. HPASMC were transfected with 2µg of plasmid over-expressing lnc536 (PUC57-lnc536), or PUC57-empty as control (n=3) for 24h followed by RNA-IP using RBM25 antibody. (B) RNA-IP of PDGF-treated HPASMCs using RBM25 antibody showed an increase in lnc536 (i) binding with RBM25. Corresponding lower levels of HOXB13 (ii) were observed. (C) Effect of knockdown of RBM25 in HPASMCs over-expressing lnc-536 on HOXB13 expression (i) and cell proliferation as assayed by MTS (ii), Cyquant (iii) and Edu (iv). HPASMCs transfected with scrambled RNA (Scr-RNA) or small hairpin against RBM25 (RBM25 shRNA). (D) Effect of knockdown of RBM25 in HPASMCs treated with PDGF on (i) HOXB13 expression and cell proliferation as measured by MTS (ii) Cyquant (iii) and Edu (iv). HPASMCs were either transfected with shRBM25 or Scr-RNA followed by treatment with and without PDGF (100ng/ml). * p<0.05, **p <0.01, *** p<0.001. The data shown is the average of n=3 experiments. Representative Edu-stained images are shown.

### Knock-down of RBM25 in HPASMCs results in the up-regulation of HOXB-13 expression leading to attenuation in cell proliferation

To further examine if lnc536 exerts its effects on HOXB13 expression and proliferation through RBM25 interactions, we knocked down RBM25 in HPASMCs over expressing lnc-536 or treated with PDGF. The co-transfection of cells with PUC57-lnc536 and shRNA against RBM25 shRNA resulted in increased expression of HOXB13 compared to lnc-536 overexpressing cells with control levels of RBM25 (PUC57-lnc536 + RBM25 shRNA vs PUC57-lnc536) or compared to cells with only knockdown of RBM25 (PUC57-lnc536 + RBM25 shRNA vs RBM25 shRNA) (Figure 3Ci). Parallel to this, proliferation was also found to attenuate in lnc536 overexpressing cells after knockdown of RBM25 (PUC57-lnc536 + RBM25 shRNA vs PUC57-lnc536) (Figure 3C ii-iv), suggesting that RBM25 is a mediator of the lnc536-HOXB13 interactions regulating cell proliferation.

Similarly, the knock-down of RBM25 in PDGF-treated HPASMCs increased the expression of HOXB13 and decreased the proliferation of cells compared to scrambled control (RBM25 shRNA +PDGF vs Scr RNA +PDGF) as shown in Figure 3D (i-iv). These results suggest that RBM25-mediated lnc536-HOXB13 interactions lead to a decrease in HOXB13 and augmentation of smooth muscle proliferation. Interestingly, RBM25 knockdown alone showed increased proliferation compared to scrambled control suggesting the multifunctional role of RBM25 in the transcriptional and post-transcriptional regulation of various proliferative and anti-proliferative genes. However, this increase in proliferation after RBM25 knockdown was attenuated in cells treated with PDGF with a corresponding increase in HOXB13 (RBM25 shRNA +PDGF vs RBM25 shRNA) (Figure 3D).

### Lnc536-RBM25 complex acts as a decoy for the Splicing Factor Proline/Glutamine Rich (SFPQ) transcriptional regulator thereby reducing HOXB13 expression

The STRING (Search Tool for the Retrieval of Interacting Genes/Proteins) analysis of pulled-down proteins by lnc-536 identified protein-protein interactions (direct physical or indirect functional correlations) of RBM25 with other 20 pulled-down proteins. The effect of these interactions on the altered transcription of HOXB13 was predicted using the RBPmap bioinformatics tool and checking for RNA motif of these RNA binding proteins (RBPs) on the HOXB13 gene. We found 4 of 20 RBM25 interacting proteins to have binding motifs on HOXB13 exons (Supplementary Figure. E2F), all with Unused ProtScore of >2. Among these 4 RBM25 interacting proteins, SFPQ (a.k.a. PSF) with the highest Unused score and Interaction confidence score has been demonstrated to act as a transcriptional regulator^29–33^.

To validate if Lnc-536 -RBM25 acts by altering the SFPQ-HOXB13 complex, we first performed RNA-IP on PDGF-treated HPASMC by using SFPQ antibody and checked the levels of RBM25 by western blot in the pulled-down complex (Figure 4A). The interaction of RBM25 with SFPQ was significantly high in PDGF-treated cells. This suggests that PDGF treatment of HPASMCs with augmented levels of lnc-536 increases interactions of RBM25 with SFPQ which in turn may be responsible for the decrease in the HOXB13 expression. This was further validated by over-expressing SFPQ using plasmid (pCMV-SFPQ) in the presence and absence of PDGF in HPASMCs. It was observed that HOXB13 expression significantly increased in cells over-expressing SFPQ (pCMV-SFPQ vs pCMV-Empty) however, the addition of PDGF attenuated the increase in HOXB13 expression (pCMV-SFPQ vs pCMV-SFPQ+ PDGF) (p<0.05) (Figure 4B).

**Figure 4.**
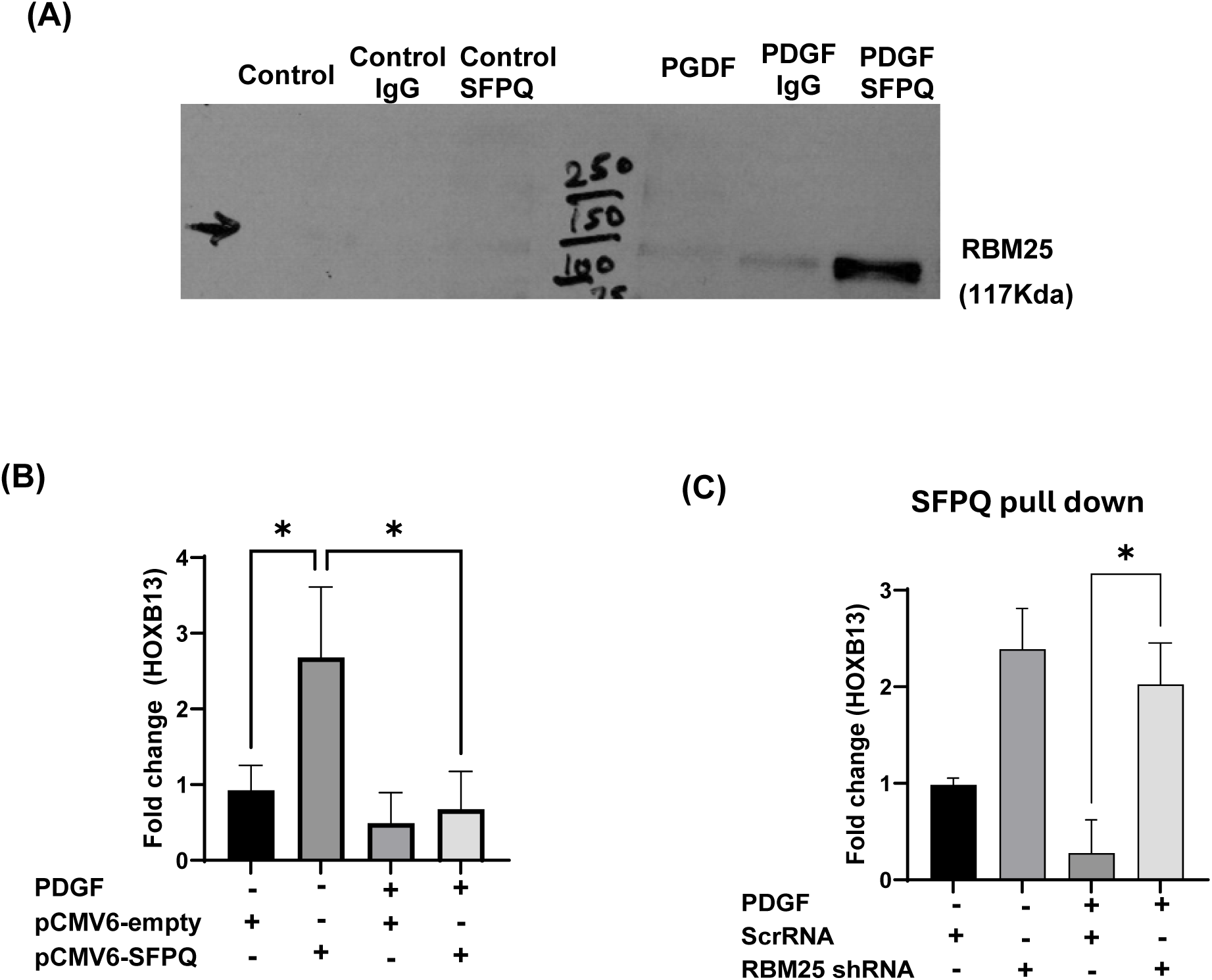
RBM25 acts as a decoy for a transcription factor: splicing factor proline/glutamine-rich (SFPQ) leading to decreased expression of HOXB13 in hyper-proliferative PDGF-treated smooth muscle cells. (A) HPASMC were treated for 24h with PDGF (100ng/ml), and IP was performed using SFPQ antibody followed by western blot analysis for RBM25. (B) Effect of SFPQ over-expression on HOXB13 levels in smooth muscle cells as assessed by qRT-PCR. HPASMCs were transfected with 2µg plasmid overexpressing SFPQ (pCMV6-SFPQ) or empty plasmid control (pCMV6-empty) for 24h followed by treated with PDGF (C) Levels of HOXB13 mRNA bound to SFPQ in HPASMCs with and without knockdown of RBM25. HPASMC transfected with RBM25 shRNA or scrRNA were treated with and without PDGF (100ng/ml) for 24h followed by RNA-IP using SFPQ antibody and HOXB13 RT-PCR analysis of pulled down RNA (n=3). * p<0.05. The data shown is the average of n=3 experiments.

To check if the PDGF-mediated increased interactions of lnc536 with RBM25 (Fig 3) and the above-mentioned interactions of RBM25 with SFPQ altered the ability of SFPQ to bind on its target HOXB13 gene promoter, we measured HOXB13 mRNA levels in complexes pulled down after immune precipitation using SFPQ antibody from HPASMC treated with and without PDGF in the presence and absence of knockdown of RBM25. As illustrated in Figure 4C, levels of HOXB13 complexed with SFPQ increased with the knockdown of RBM25 in the presence of PDGF treatment. Overall, all these findings suggest that SFPQ is involved in binding to HOXB13 promoter and activates its expression, however, PDGF-mediated increased interactions of lnc536 and RBM25 sequester away transcription factor SFPQ from the promoter region of HOXB13 and this in turn reduces the HOXB13 expression.

### Lnc-536 /HOXB13 axis in hyperproliferative smooth muscle cells from IPAH patients

Next, we examined the lnc536/RBM25 /HoxB13 interactions in hyperproliferative, apoptotic-resistant smooth muscle cells from iPAH patients consistently showed augmented expression of lnc536 in iPAH-PASMCs than in PASMCs from healthy donors (Figure 5Ai), whereas no change in the lnc536 expression was observed in cells from familial PAH (FPAH) when compared with healthy donors (Figure 5Aiii). In parallel, HOXB13 was found to have a downward trend in PASMCs from iPAH-compared to healthy cells (Figure 5Aii) while no change was observed in FPAH cells (Figure 5Aiv). The knockdown in the expression lnc-536 in IPAH cells increased the levels of HOXB13 with an associated decrease in cell proliferation (LNA-Scr-IPAH vs LNA-AS536-IPAH) (Figure 5B i-ii).

**Figure 5.**
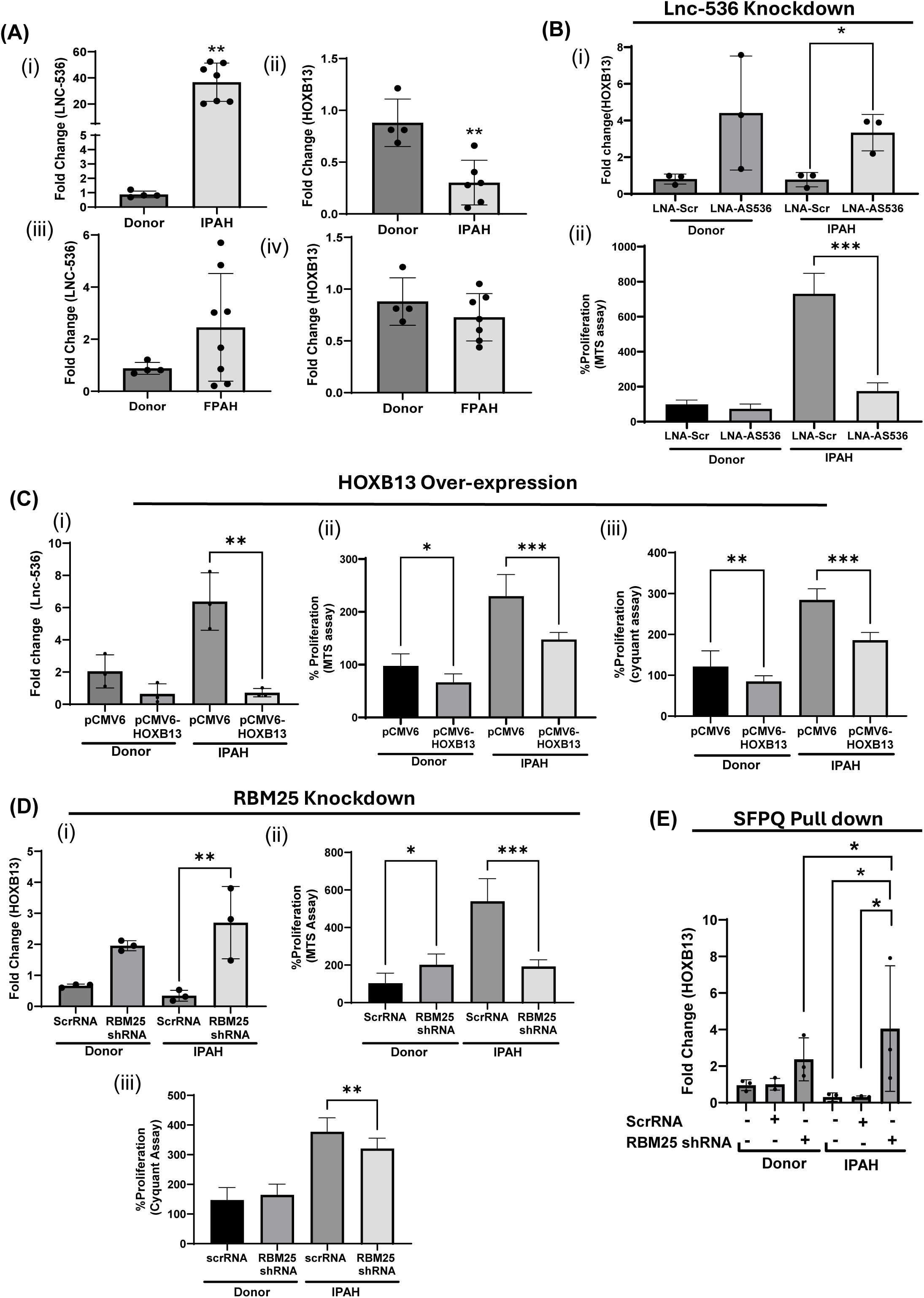
(A) Idiopathic pulmonary arterial smooth muscle cells (IPAH) show higher expression of lncRNA536 (i) and decreased expression of HOXB13 (ii) whereas no change in the expression of lncRNA536 (iii) and (iv) HOXB13 was observed in familial pulmonary arterial smooth muscle cells (FPAH). (B) Effect of lnc536 knockdown on (i) HOXB13 expression and cell proliferation as assayed by (ii) MTS. Cells obtained from IPAH and healthy donors were transfected with LNA-GapmeR against lnc536 (lnc536-AS1,40nM) or LNA-GapmeR scrambled control (n=3) for 24h.*p<0.05, *** p<0.001. (C) To study the effect of HOXB13 over-expression on IPAH cells (i) LNC536 expression (n=3) and cell proliferation as assayed using MTS (ii) and Cyquant (iii). *p<0.05, **p <0.01 and *** p<0.001. IPAH cells were treated with HOXB13 over-expressing plasmid (2ug) for 24h. (D) Effect of knockdown of RBM25 in IPAH cells on HOXB13 expression (i), cell proliferation assays as assessed by MTS (ii) and Cyquant (iii). IPAH cells were transfected with ScrRNA (control) and RBM25 shRNA (knockdown) (n=3) for 24h, (E) Effect of knockdown of RBM25 in IPAH cells and RNA-IP using SFPQ antibody on HOXB13 expression (n=3). *p<0.05, **p <0.01, *** p<0.001.

Meanwhile, the transfection of IPAH-PASMCs with plasmid over-expressing HOXB13 (pCMV6-HOXB13) resulted in a feed-forward decrease in the expression of lnc536 (pCMV6 empty+ IPAH vs pCMV-HOXB13+ IPAH) and the decrease in proliferation as shown in Figure 5C (i-iii). Further, to validate the specific interaction between RBM25 and lnc536, we knocked down the expression of RBM25 in IPAH-PASMCs using RBM25 shRNA. As expected, the knockdown of RBM25 in IPAH cells led to a significant increase in the expression of HOXB13 (ScrRNA + IPAH vs RBM25shRNA+IPAH) along with a decrease in proliferation (Figure 5D i-iii).

Further, the RNA-IP analysis using SFPQ antibody found higher levels of HOXB13 bound to SFPQ in IPAH cells after the knockdown of RBM25 (ScrRNA +IPAH vs RBM25shRNA+ IPAH) suggesting the inhibitory role of RBM25 in the interactions between SFPQ and HOXB13 promoter (Figure 5E). All these findings confirm that LNC536 contributes to the RBM25-mediated remodeling of the SFPQ-HOXB13 complex in the hyperproliferative smooth muscle cells derived from IPAH patients.

### Knockdown of Lnc-536 attenuates smooth muscle hyperplasia and pulmonary vascular dysfunction in response to the dual hit of HIV and cocaine

We also observed a decrease in the expression of lnc536 in cocaine and HIV-Tat (C+T) treated hyperproliferative HPASMCs transfected with plasmid over-expressing HOXB13 (pCMV6 empty C+T vs pCMV6 HOXB13 C+T) with a parallel decrease in C+T mediated cell proliferation (Figure 6A (i-iii). Further, we observed a significant increase in the expression of HOXB13 along with a decrease in proliferation in C+T treated cells after knock-down of RBM25 from HPASMCs (ScrRNA C+T vs RBM25shRNA C+T) (Figure 6Bi-iii) suggesting RBM25 as a mediator of lnc536-HOXB13 interaction in cocaine and Tat-treated hyperproliferative smooth muscle cells.

**Figure 6.**
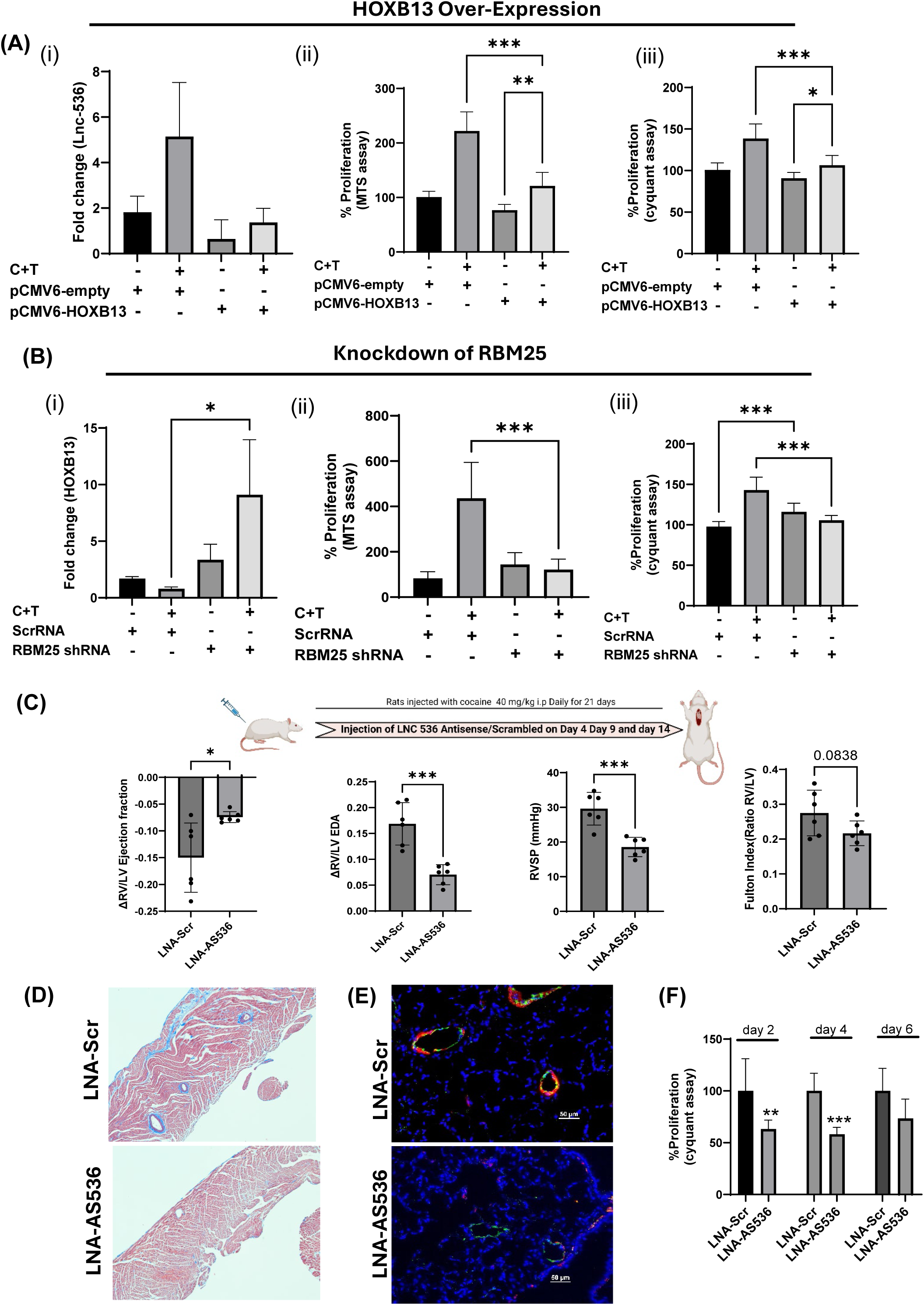
Involvement of lnc-536/RBM25 /HOXB13 axis in HIV and cocaine-mediated smooth muscle hyperplasia. (A) Effect of cocaine and HIV protein Tat (C+T) treatment on lnc536 expression (i) and cell proliferation as measured by MTS (ii) and Cyquant (iii) in HPASMCs over expressing HOXB13. HPASMCs were transfected with either plasmid over-expressing HOXB13 (pCMV6-HOXB13) or empty plasmid (pCMV6-empty) for 24h followed by treatment with cocaine (C) (1 µM and Tat (T) (25 ng/mL) for 48h. (B) Effect of knockdown of RBM25 on HOXB13 expression (i), cell proliferation as measured by MTS (ii) and Cyquant (iii). Cells were transfected with shRBM25 or scrRNA for 24h followed by treatment with C+T for 48h. * p<0.05, **p <0.01, *** p<0.001. (C) Effect of knockdown of lnc536 in HIV-transgenic cocaine treated PAH rats on normalized Δ EV/LV Ejection fraction (EF), ΔRV/LV end-diastolic area (EDA), RVSP, and Fulton index. HIV-Tg male Fisher rats (n = 6–8/group exposed to cocaine daily for 21 days were treated with LNA-scrambled GapmeR control (LNA-Scr) or LNA GapmeR against lnc536 (LNA-AS 536) on days 4, 9, and 14. (D) Representative images of trichrome stained right ventricles and (E) pulmonary vascular remodeling as represented by alpha-smooth muscle actin (red) and vWF ( green) immuno-fluorescence staining in lungs Scale bars, 100 mm. (F) Cyquant proliferation analysis of PASMCs isolated from rats grown under no serum conditions for 2, 4 or 6 days *p <0.01, *** p<0.001.

Previously we reported an increase in mean pulmonary arterial pressure (mPAP) and right ventricle systolic pressure (RVSP) in HIV Tg rats exposed to cocaine ^34^ and this was associated with enhanced pulmonary vascular remodeling and smooth muscle proliferation. Now to check the *in-vivo* role of lnc-536 in response to the dual hit of cocaine and HIV-1 in the development of PAH, we injected LNA-AS-536 GapmeRs in HIV-Tg rats exposed to cocaine. As shown in Figure 6C, treatment of HIV-Tg rats with AS536 resulted in a significant increase in ΔRV-EF and a significant decrease in ΔRV-EDA and RVSP when compared with HIV-Tg rats treated with LNA-Scr group rats with no change in mean BP, heart rate or cardiac output between the groups (Supplementary Figure E3A). Although the downward trends in the Fulton index on treatment with AS536 were found to be not statistically significant, trichrome staining of right ventricles confirmed the protective effect of AS-536 as shown by the representative micrograph (Figure 6D) with less fibrosis in the RV from LNA-AS536 rat when compared with LNA-Scr rat. In addition, lack or reduced smooth muscle thickening was observed in small and medium-sized vessels in HIV Tg cocaine-exposed rats treated with LNA-AS536 (Figure 6E). This corresponded with the reduction in the lnc536 and augmentation in the HOXB13 expression in the PASMC isolated from HIV-Tg rats treated with LNA-AS536 compared to control rats exposed to LNA-Scr GapmeRs (Supplementary Figure E3B). In addition, treatment with LNA-AS536 markedly attenuated the proliferation of PASMCs in these rats as validated by Cyquant (Figure 6F), MTS, and Edu staining assays (Supplementary Figure E3C-D).

## DISCUSSION

Recently, long non-coding RNAs have emerged as important regulators of diverse biological processes including cell proliferation and apoptosis ^35,36^. Our findings suggest that the hyperproliferative phenotype of IPAH patient-derived PASMCs and PDGF- or even HIV protein and cocaine stimulator-treated cells is regulated by elevated levels of lnc536 via downregulating the expression of the anti-proliferative HOXB13. In vivo, knockdown of lnc536 in Sugen-Hypoxia and HIV Tg PAH rats using AS-536 LNA GapmeRs reversed the enhanced RVSP and pulmonary vascular remodeling. The RNA-pull-down experiments and RNA-IP analysis revealed that lnc536 acts as a decoy for RBM25 and this disrupts the binding of the transcription regulator and SFPQ to the HOXB13 promoter leading to a downregulation of HOXB13 expression and a reduction in PASMC proliferation (Figure 7).

**Figure 7:**
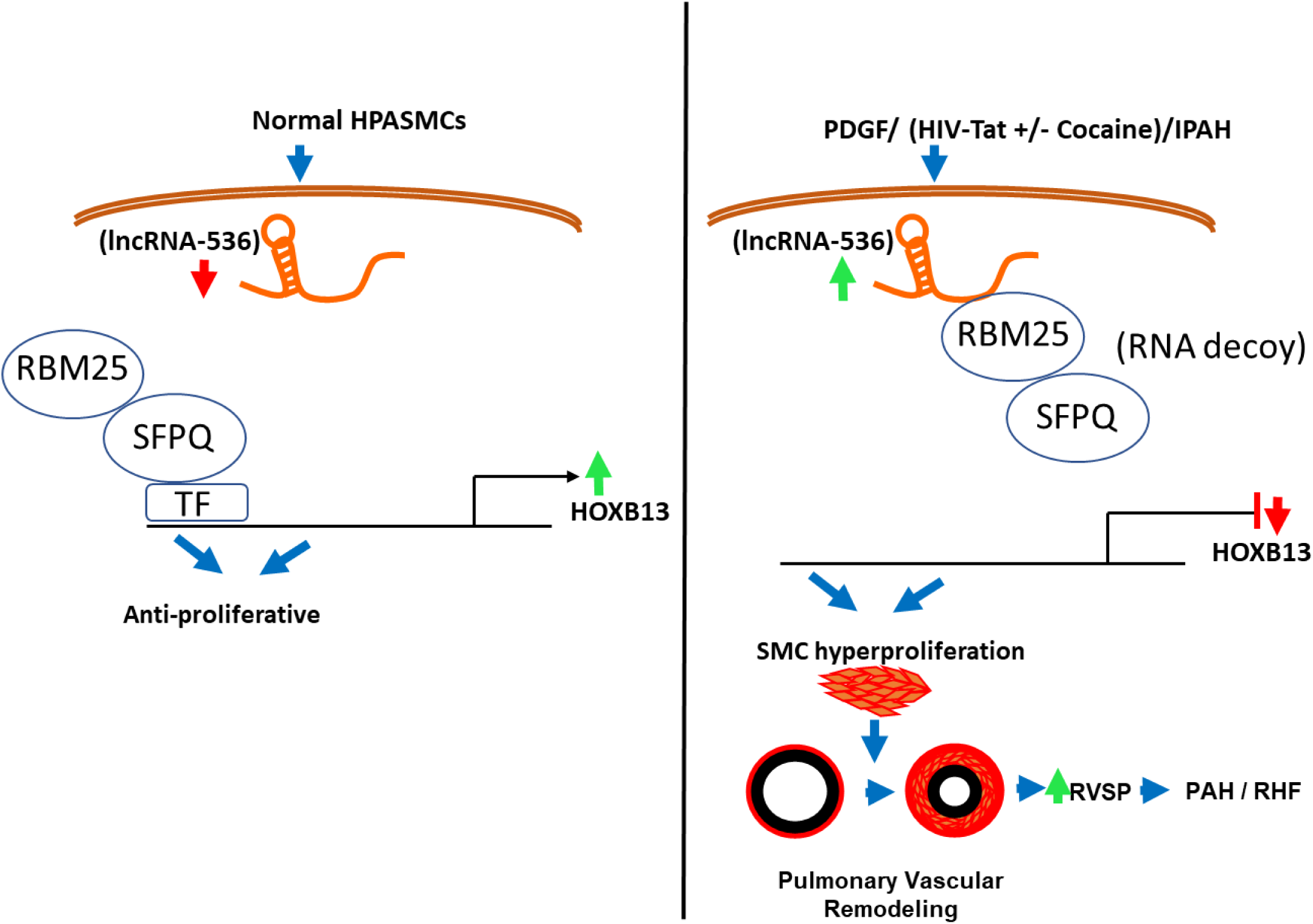
Schematic showing the lnc536 mediated regulation of hyperproliferative phenotype of pulmonary arterial smooth muscle cells in PAH. Lnc-536 is up-regulated in IPAH patient-derived PASMCs and in response to PDGF mitogen or HIV protein and cocaine stimulant, Lnc-536 acts as a decoy for an RNA binding protein, RBM25, which results in the sequestering of transcription regulator: SFPQ. This prevents SFPQ from binding to the promoter region of the anti-proliferative HOXB13 gene thereby decreasing its expression and augmenting the proliferation of PASMCs.

Studies have shown that dysregulation of lncRNAs is linked to various human diseases, including cancers and neurological disorders ^8,9^, but their role in pulmonary hypertension is not well-documented. LncRNA H19 was found to be increased in the monocrotaline (MCT)-induced animal model of PAH ^37^. Pullamsetti lab showed increased levels of lncRNA TYKRIL in the hypoxia-exposed smooth muscle cells and in IPAH patients ^38^. In addition, lncRNA Gas5 was reduced with hypoxia-induced PAH in rats ^39^ while hypoxia-stimulated human PASMCs showed reduced levels of Gas5 and lncRNA CASC2 ^39,40^. In a few cases, lncRNA-mediated regulatory roles and mechanisms in PAH remain contradictory. For example, MEG3 was found to be downregulated in PASMCs derived from patients with PAH ^41^) whereas it was upregulated in hypoxia-induced PAH in a mice model ^42^. We here identified and investigated the role of the novel lncRNA lnc536 in PAH. Notably, our results from IPAH patient-derived PASMCs and HPASMCs treated with PDGF or with combined treatment of cocaine and HIV-Tat demonstrated that lnc536 was broadly elevated in all pro-proliferative settings.

Increased levels of lnc-536 in PDGF-treated or IPAH PASMCs were found to downregulate the HOXB13 expression. HOXB13 is the last member of the HOXB cluster which acts as a tumor suppressor and the altered expression of HOXB13 has been reported in many cancers^43–46^. Loss of HOXB13 in colorectal tumor cells has been associated with increased proliferation^45,46^. The tumor suppressor HOXB13 is reported to inhibit pro-proliferative Wnt/β-catenin signaling in prostate cancer^45^ The literature also suggests that the forced expression of HOXB13 in colorectal tumor cells directly suppresses the transactivation of T cell factor 4 (TCF4) expression leading to inhibition of β -catenin/TCF4 signaling and cell growth ^46^. Furthermore, the inhibitory effects of HOXB13 on cell cycle progression via proteasomal degradation of cyclin D1 have been shown in prostate cancer ^43^.

The RNA-pull-down assay and RNA-IP analysis suggested that lnc536 potentially interacts with RBM25 resulting in the downregulation of HOXB13 expression and augmentation of HPASMC proliferation. Knockdown of RBM25 in PDGF-treated and IPAH HPASMCs or in overexpressing lnc536 resulted in an increase in HOXB13 expression along with an attenuation in HPASMC proliferation. To the best of our knowledge, LNC536 is the first lncRNA to be observed to regulate the RBM25/HOXB13 axis. RBM25, a highly conserved putative splicing factor is a potential tumor suppressor^28^ and is known to interact with several splicing factors. RBM25 was previously found to be elevated in human heart failure tissue and is known to promote the generation of non-functional splicing variants of SCN5a in heart failure and pulmonary hypertension ^47^. This RBM25-mediated abnormal SCN5A mRNA splicing is known to reduce Na+ channel current which results in sudden death ^48^. In addition, RBM25 is reported to inhibit proliferation by promoting the formation of active isoform of BIN1 (Myc-inhibitor) ^22^. It also promotes apoptosis by increasing the splicing of anti-apoptotic BCL-XL to pro-apoptotic BCL-XS isoform^49^. Therefore, there is a possibility that lnc-536 may also be involved in trans-regulating the BIN1 and BCL-XL/S splicing or regulating SCN5A splice variants by acting as a decoy or guide to RBM25, resulting in the activation of pro-proliferative/anti-apoptotic signaling.

In examining the molecular mechanisms underlying the lncRNA-RBM25 regulation of HOXB13 expression, our findings demonstrate that lncRNA-536 acts as a decoy for RBM25 which in turn sequesters SFPQ making it unavailable to bind to the promoter region of anti-proliferative HOXB13 and induce its expression; thereby promoting proliferative phenotype of pulmonary smooth muscle cells associated with PAH development. Splicing Factor Proline and Glutamine Rich (SFPQ) is a widely distributed nuclear protein that interacts with dsDNA at the promoter site of several genes and regulates their transcription ^50^. Furthermore, SFPQ, commonly referred to as PSF (PTB-associated splicing factor) functions as a splicing-related factor in mRNA processing and is crucial for the repair of double-strand break DNA damage ^51–53^. SFPQ can act as a transcription activator or repressor. depending on the cellular environment. SFPQ mediates transcriptional activation of adenosine deaminase B2 (ADARB2) by binding to its receptor whereas reduces the STAT6-dependent transcription by recruitment of co-repressor SIN3A and histone deacetylases (HDACs) or histone methyltransferases ^54,55^. It has been documented that SFPQ interacts with lncRNAs, such as NEAT1 which is an essential factor for the establishment of subnuclear ’paraspeckle’ and is involved in the regulation of SFPQ-mediated splicing regulation. Additionally, lncRNA MALAT1 interacts with SFPQ to impede the association of SFPQ and PTBP2 proteins, which then mediates the release of oncogene PTBP2 from the SFPQ/PTBP2 complex, resulting in tumor growth and metastasis ^30^. Similarly, our current findings suggest that lnc536 impacts SFPQ’s regulatory role on HOXB13 expression by sequestering SFPQ /RBM25 complex away from the HOXB13 promoter. In the context of pulmonary hypertension, SFPQ has been reported to attenuate the transcription of CD40 involved in proliferation, migration, and pro-inflammatory activity of rat pulmonary artery adventitial fibroblasts by forming a complex with histone deacetylase 1 (HDAC1)^56^.

Although SFPQ displayed the highest Unused ProtScore and Interaction Confidence Score among the 4 RBM25 interacting proteins that had motifs for binding HOXB13, twenty additional proteins, including EIF5b, PRPF40A that were pulled down by lnc-536 may also be involved in regulating smooth muscle proliferation by targeting different downstream anti- or pro-proliferative molecules. Further, we didn’t analyze lncRNA expression using the precision-cut lung sections from PAH patients, however, the analysis using HPASMCs from IPAH patients and our in-vivo studies using rat PAH models, considering the conservation of lncRNA in rat species, add to the significance and translational aspect of the study.

In conclusion, we report that lnc536 contributes significantly to the pulmonary vessel remodeling associated with PAH by promoting the hyperproliferation of pulmonary arterial smooth muscle cells in both in-vitro and in-vivo conditions and thus represent a novel therapeutic target for pulmonary hypertension in general. To the best of our knowledge, we identified for the first time that Lnc-536 acts as a decoy for an RNA binding protein, RBM25 which in turn sequesters transcription regulator SFPQ. This prevents SFPQ from binding to the promoter region of the anti-proliferative HOXB13 gene thereby decreasing its expression and augmenting proliferation of PASMCs during PAH development. Better insight in understanding the role of lncRNAs in the pathogenesis of pulmonary vascular remodeling associated with cardio-pulmonary complications is expected to is expected to have significant translational impact by opening new avenues for potential therapeutic interventions.

## ACKNOWLEDGMENTS

The funds to carry out the study were provided by National Institute of Health (NIH) grants R01DA040392 and, R01 HL152832 awarded to N.K.D. IPAH cells were provided by PHBI under the Pulmonary Hypertension Breakthrough Initiative (PHBI). Funding for the PHBI was provided under an NHLBI R24 grant, #R24HL123767, and by the Cardiovascular Medical Research and Education Fund (CMREF).

## Author Contributions

All cell-culture experiments were performed by A.M, and L.C. A.K and L.C performed animal experiments. A.M, A.K, and N.K.D. analyzed and interpreted the data. Statistical analysis was performed by A.M, and A.K. A.M and N.K.D. contributed to writing the manuscript. A.M and N.K.D. designed experiments and N.K.D conceptualized and supervised the research. All authors read the manuscript and approved the study.

